# CRISPulator: a discrete simulation tool for pooled genetic screens

**DOI:** 10.1101/119131

**Authors:** Tamas Nagy, Martin Kampmann

## Abstract

The rapid adoption of CRISPR technology has enabled biomedical researchers to conduct CRISPR-based genetic screens in a pooled format. The quality of results from such screens is heavily dependent on optimal screen design, which also affects cost and scalability. We present CRISPulator, a computational tool that simulates the impact of screen parameters on the robustness of screen results, thereby enabling users to build intuition and insights that will inform their experimental strategy. We illustrate its power by deriving non-obvious rules for optimal screen design.

## Background

Genetic screening is a powerful discovery tool in biology that provides an important functional complement to observational genomics. Until recently, screens in mammalian cells were implemented primarily based on RNA interference (RNAi) technology. Inherent off-target effects of RNAi screens present a major challenge [1]. In principle, this problem can be overcome using optimized ultra-complex RNAi libraries [2, 3], but the resulting scale of the experiment in terms of the number of cells required to be screened can be prohibitive for some applications, such as screens in primary cells or mouse xenografts.

Recently, several platforms for mammalian cell screens have been implemented based on CRISPR technology [4]. CRISPR nuclease (CRISPRn) screens [5, 6] perturb gene function by targeting Cas9 nuclease programmed by a single guide RNA (sgRNA) to a genomic site inside the coding region of a gene of interest, followed by error-prone repair through the cellular non-homologous end-joining pathway. CRISPR interference (CRISPRi) and CRISPR activation (CRISPRa) screens [7] repress or activate the transcription of genes by exploiting a catalytically dead Cas9 to recruit transcriptional repressors or activators to their transcription start sites, as directed by sgRNAs.

CRISPRn and CRISPRi have vastly reduced off-target effects compared with RNAi, and thus overcome a major challenge of RNAi-based screens. However, other challenges to successful screening [1] remain. The majority of CRISPRi and CRISPRn screens have been carried out as pooled screens with lentiviral sgRNA libraries. While this pooled approach has enabled rapid generation and screening of complex libraries, successful implementation of pooled screens requires careful choices of experimental parameters. Choices for many of these parameters represent a trade-off between optimal results and cost.

## Results

Here, we present a computational tool, termed CRISPulator, which simulates how experimental parameters will affect the detection of different types of gene phenotypes in pooled CRISPR-based screens. CRISPulator is freely available online (http://crispulator.ucsf.edu) to enable researchers to develop an intuition for the impact of experimental parameters on pooled screening results, and to optimize the design of pooled screens for specific applications. It simulates all steps of pooled screens, as visualized in **Fig. 1** and described in more detail in the Methods.

**Figure 1.**
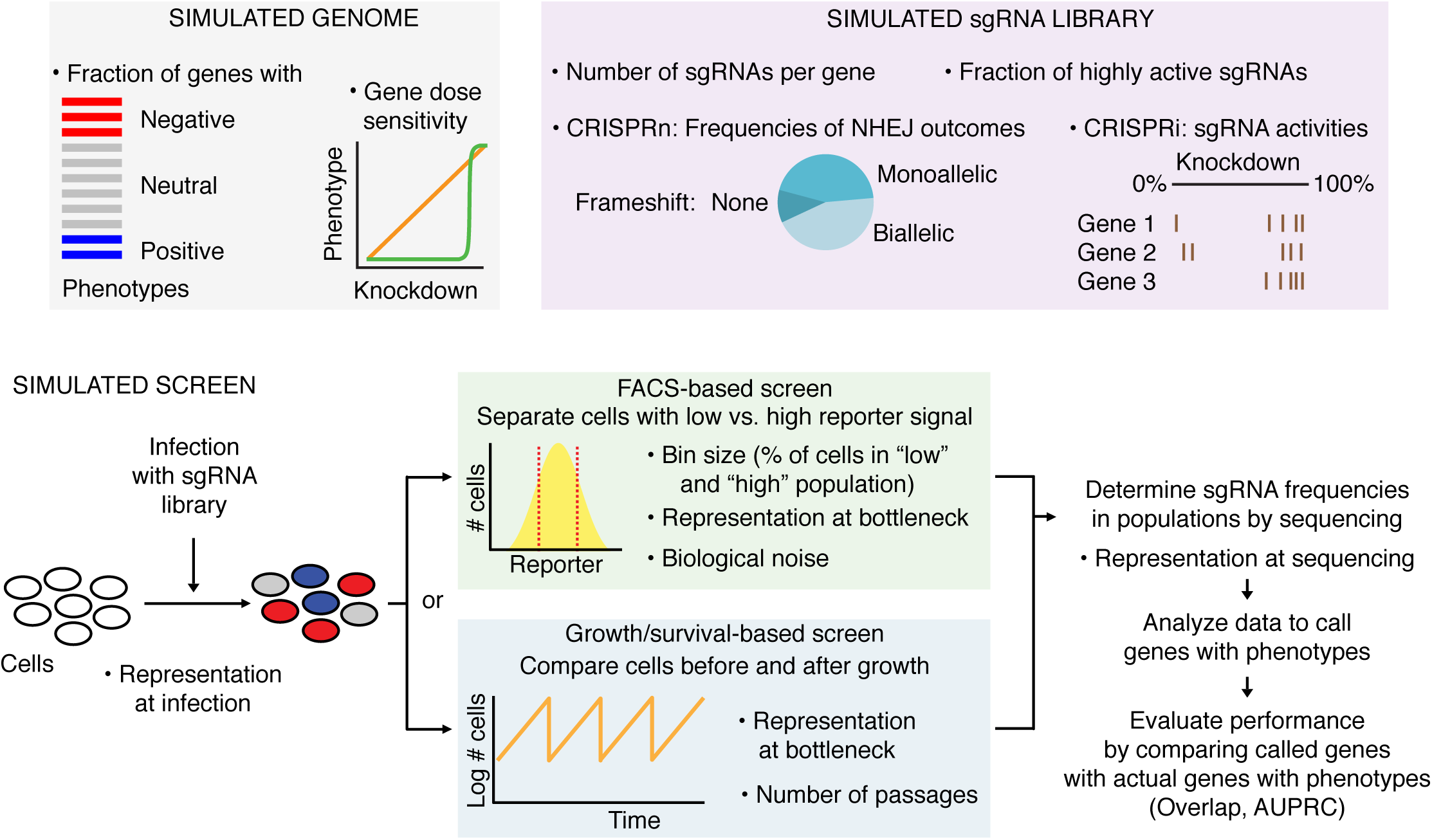
CRISPulator simulates pooled genetic screens to evaluate the effect of experimental parameters on screen performance. Overview of simulation steps: Parameters listed with bullet points can be varied to examine consequences on the performance of the screen, which is evaluated as the detection of genes with phenotypes (quantified as overlap or area under the precision-recall curve, AUPRC). Details are given in the text and Methods.

Briefly, a theoretical genome is generated in which genes are assigned quantitative phenotypes (**Fig. 2**). Independently, the quantitative relationship between gene knockdown level and resulting phenotype is defined for each gene (**Fig. 3**). Next, a sgRNA library targeting this genome is defined. Each gene is targeted by a number of independent sgRNAs. The technical performance of each sgRNA is randomly assigned based on a user-defined distribution of CRISPRn or CRISPRi sgRNA activities (**Fig. 4**), and the initial frequency distribution is specified (**Fig. 5**).

**Figure 2.**
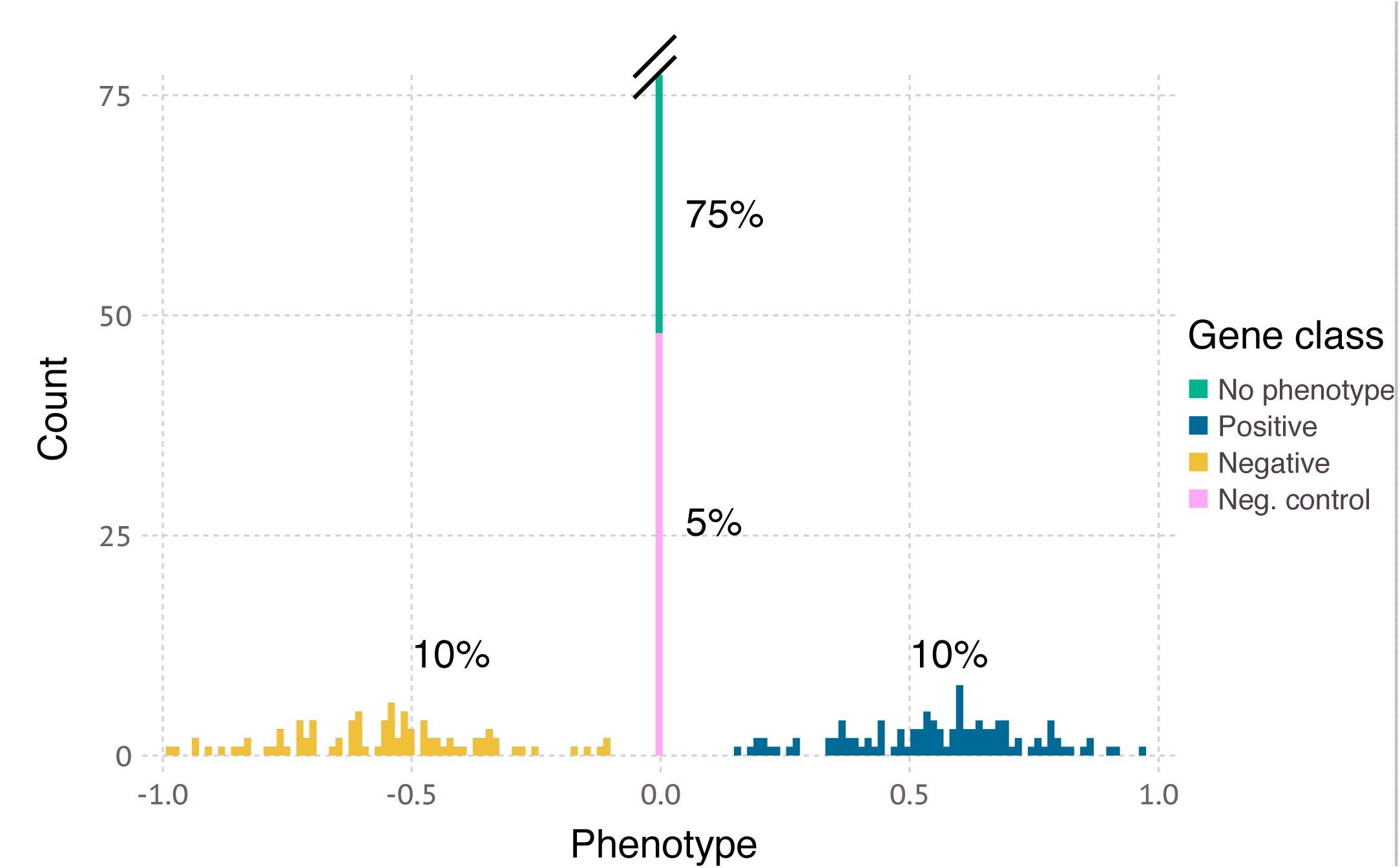
Phenotype distribution in the simulated genome. A typical distribution is shown, which includes 75% of genes without phenotype (green), 5% of negative control genes (pink), 10% of genes with a positive phenotype (blue), and 10% of genes with a negative phenotype (yellow).

**Figure 3.**
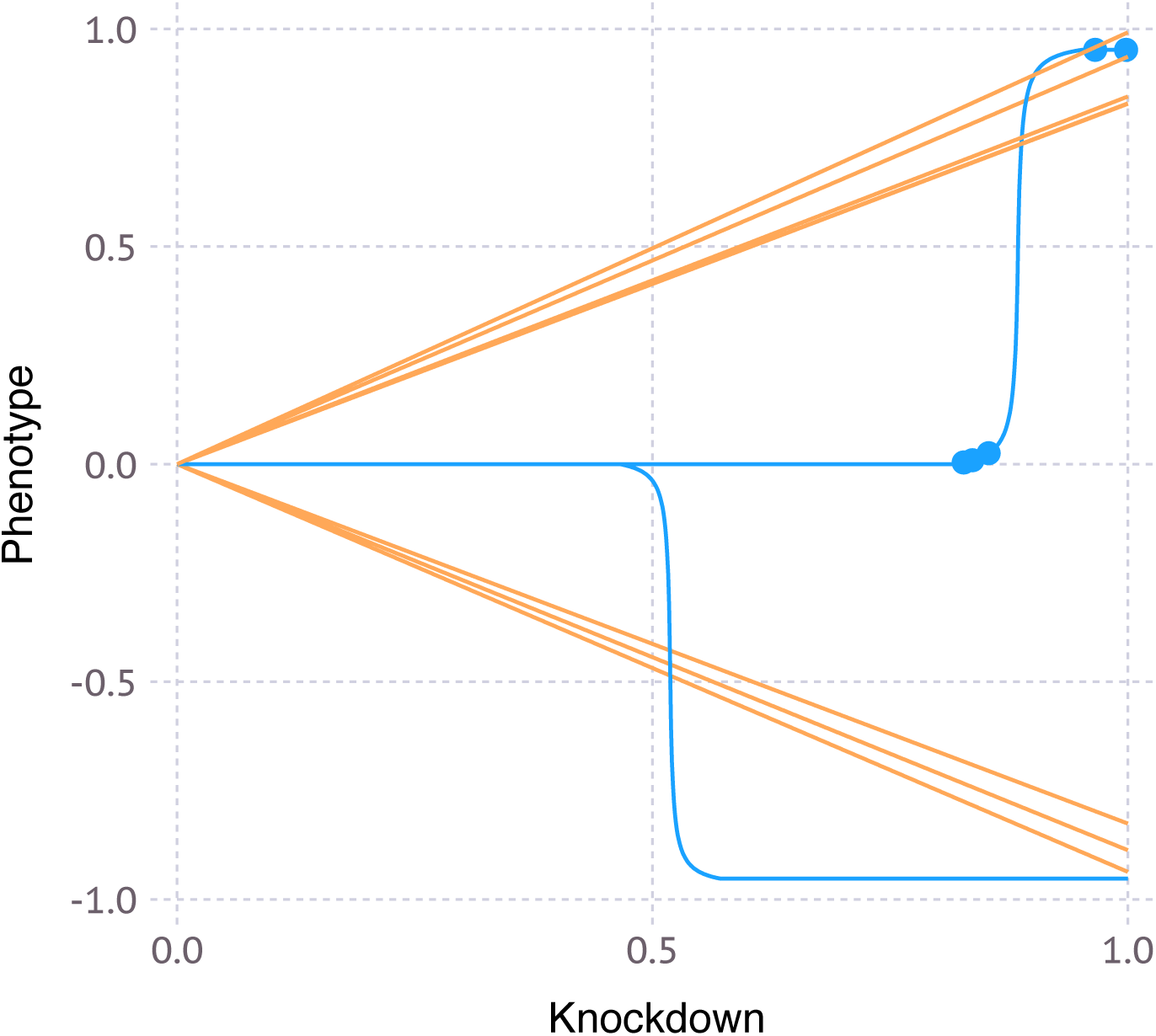
Relationship between gene knockdown level and resulting phenotype. This relationship is defined for each gene, and represents either a linear function (orange graphs) or a sigmoidal function (blue lines), as defined in the Online Methods.

**Figure 4.**
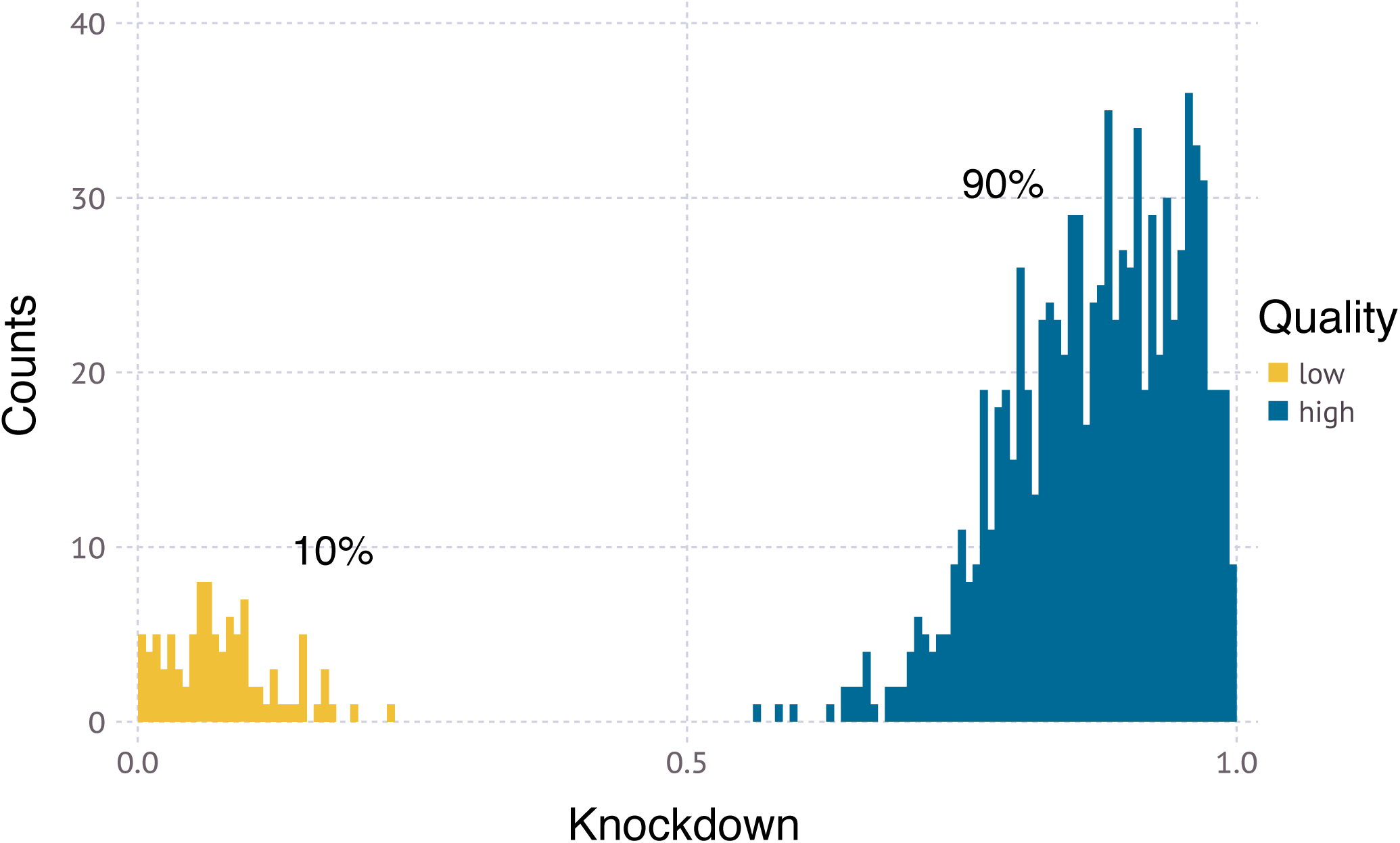
CRISPRi sgRNA activity distribution. An example of a typical distribution for 1000 guides is shown.

**Figure 5.**
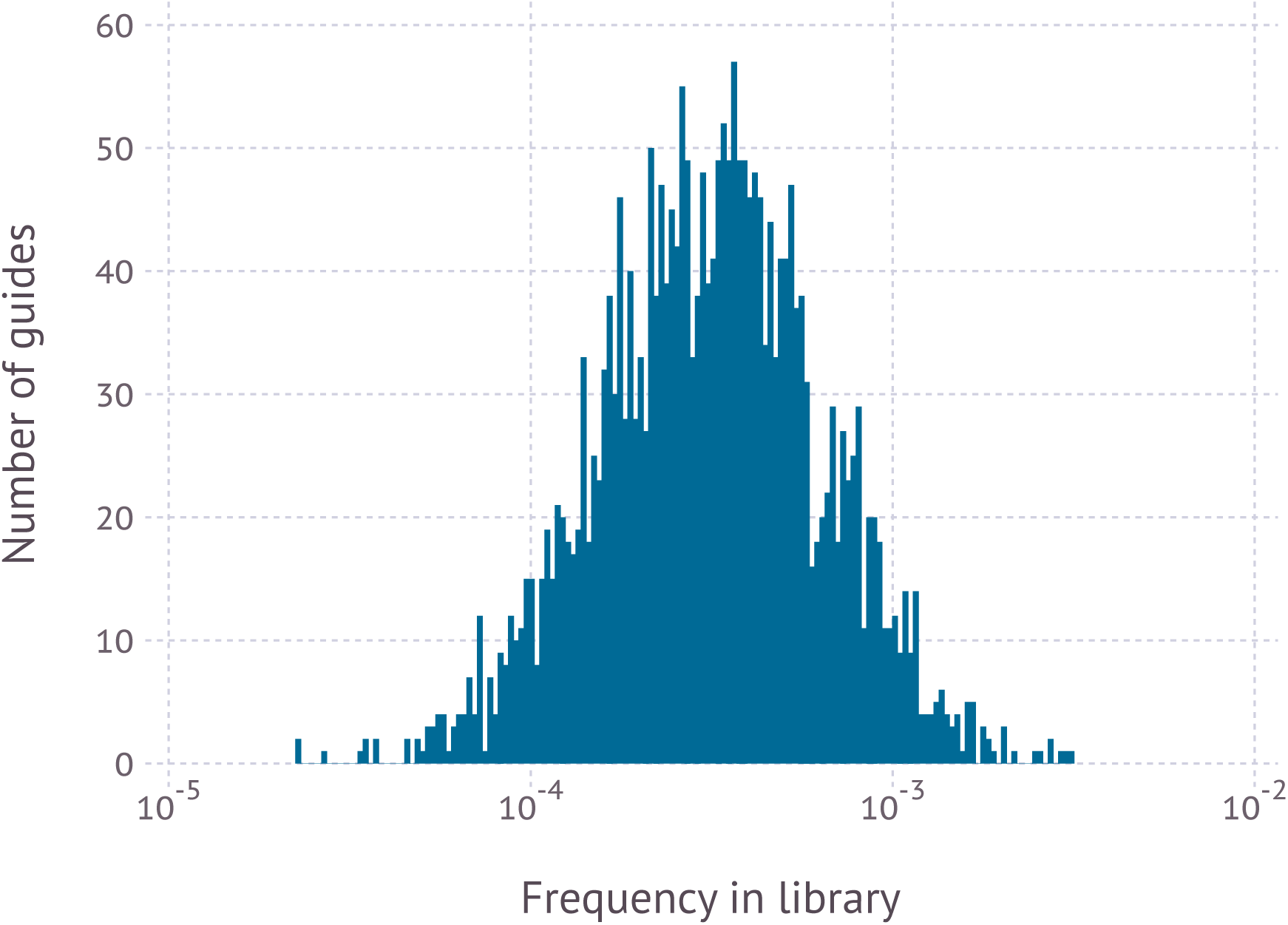
Initial frequency distribution of sgRNAs. An example of a typical distribution is shown.

Simulation of the screen itself discretely models infection of cells with the pooled sgRNA library, phenotypic selection of cells and quantification of sgRNA frequencies in selected cell populations by next-generation sequencing. Based on the resulting data, hit genes are called (**Fig. 6**) using our previously described quantitative framework [3], as detailed in the Online Methods. The performance of the screen with a specific set of experimental parameters is evaluated by comparing the called hit genes to the actual genes with phenotypes defined by the theoretical genome. It is quantified either as overlap of the list of top called hits with the actual list of top hits, or as area under the precision-recall curve (AUPRC), a metric commonly used in machine learning (**Fig. 7**).

**Figure 6.**
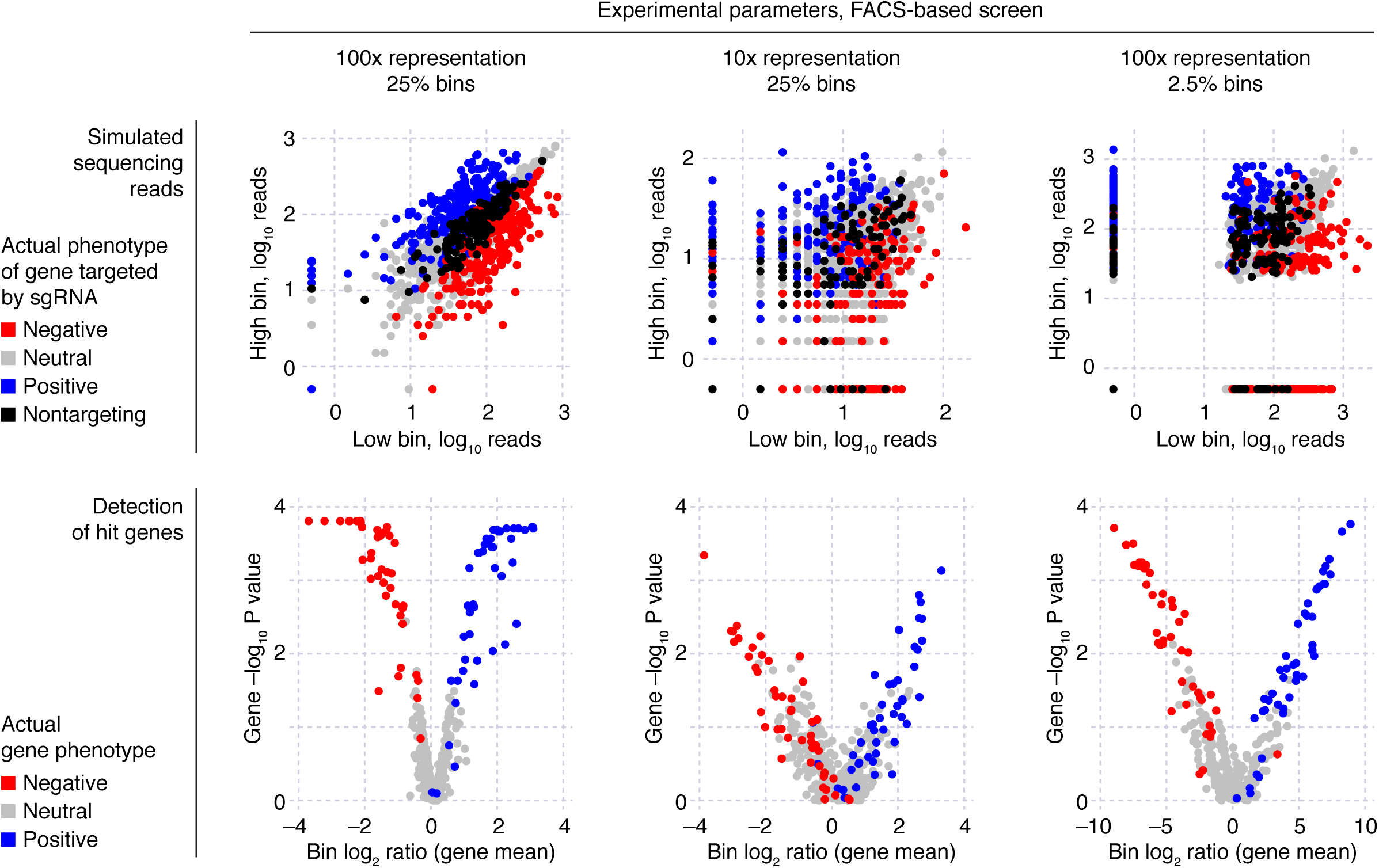
Sample results from a CRISPulator simulation of a FACS-based screen. Top row: Each point represents and individual sgRNA, plotting its read numbers in the simulated deep sequencing run for the “low reporter signal” bin and the “high reporter signal” bin. sgRNAs are color-coded to indicate whether they target a gene with a positive phenotype (knockdown increases reporter signal, blue), a gene with a negative phenotype (knockdown decreases reporter signal, red), a gene without phenotype (grey), or whether they are non-targeting control sgRNAs (black). Bottom row: Based on the observed sgRNA phenotypes, gene phenotypes are calculated (mean log_2_ ratio of read frequencies in “high” over “low” bins), and a gene P value is calculated to express statistical significance of deviation from wild-type. These are visualized in volcano plots in which each dot represents a gene. Genes are color-coded to indicate the actual phenotype: positive, blue; negative, red; no phenotype, grey.

**Figure 7.**
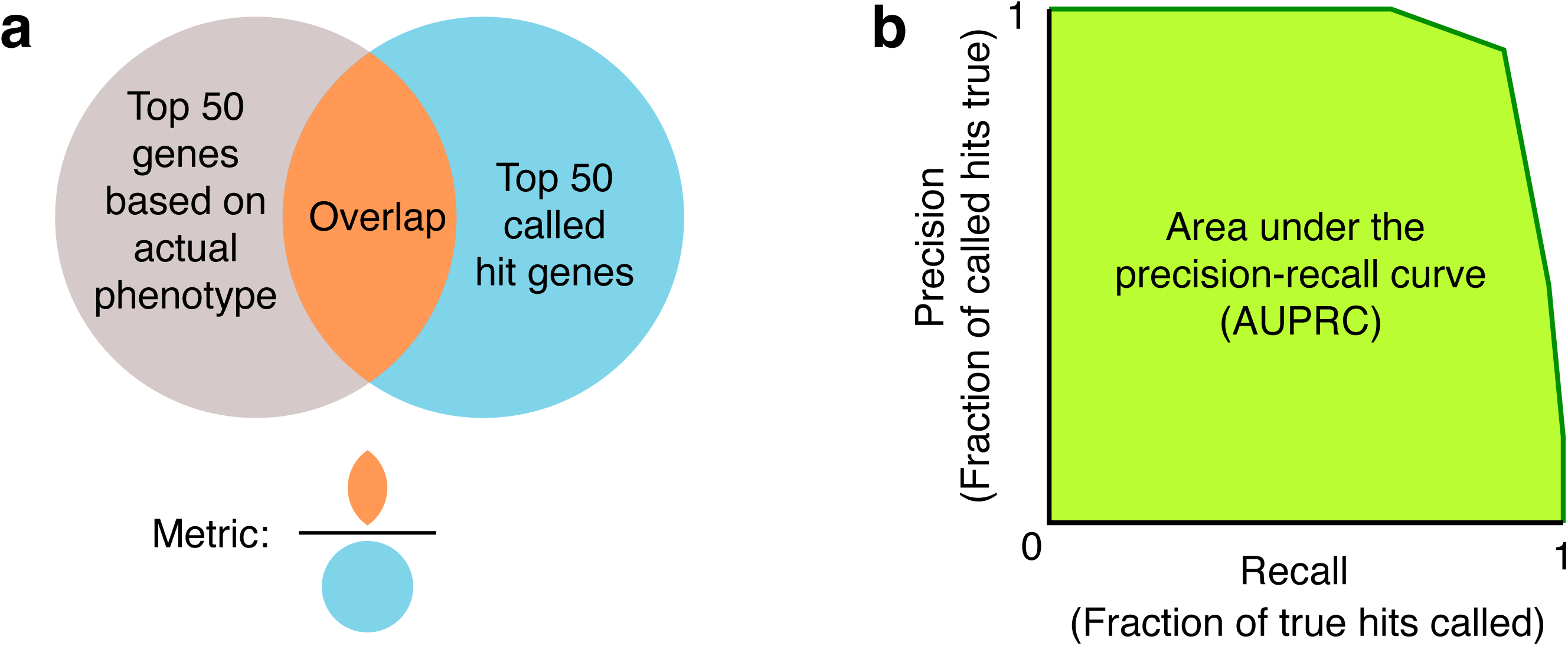
Metrics to evaluate screen performance. (**a**) “Venn diagram” overlap between the 50 genes with the strongest actual phenotypes, and the top 50 hit genes called based on the screen results – expressed as the ratio of the number of genes in the overlap over the number of called top hit genes, i.e. 50. (**b**) Area under the precision-recall curve (AUPRC).

A central consideration for all pooled screens is the number of cells used relative to the number of different sgRNAs in the library. We refer to this parameter as representation, and distinguish representation at the time of infection, representation at times during phenotypic selection, and – by extension – representation at the sequencing stage (where it is defined as the number of sequencing reads relative to the relative to the number of different sgRNAs). From first principles, higher representation is desirable to reduce Poisson sampling noise (“jackpot effects”); in practical terms, higher representation is also more costly. A major application of CRISPulator is the exploration of parameters to guide the choice of suitable representation at each step of the screen to enable researchers to strike the desired balance between screening cost and performance.

CRISPulator implements two distinct strategies for phenotypic selection. In fluorescence-activated cells sorting (FACS)-based screens, cell populations are separated based on a fluorescent reporter signal that is a function of the phenotype. We [8] and others [9] have successfully implemented such screens by isolating and comparing cell populations with the highest and the lowest reporter levels. More commonly, pooled screens are conducted to detect genes with growth or survival phenotypes [5-7] by comparing cell populations at an early time point with cells grown in the absence or presence of selective pressures, such as drugs or toxins.

We first asked how representation at the infection, selection and sequencing stages affects FACS- and growth-based screens (**Fig. 8**). The performance of FACS-based screens was most sensitive to the representation at the selection bottleneck, and least sensitive to representation at the infection stage, highlighting the importance of collecting a sufficient number of cells for each population during FACS sorting, ideally more than 100-fold the number of different library elements. By contrast, the performance of growth-based screens was similarly sensitive to representation at all stages.

**Figure 8.**
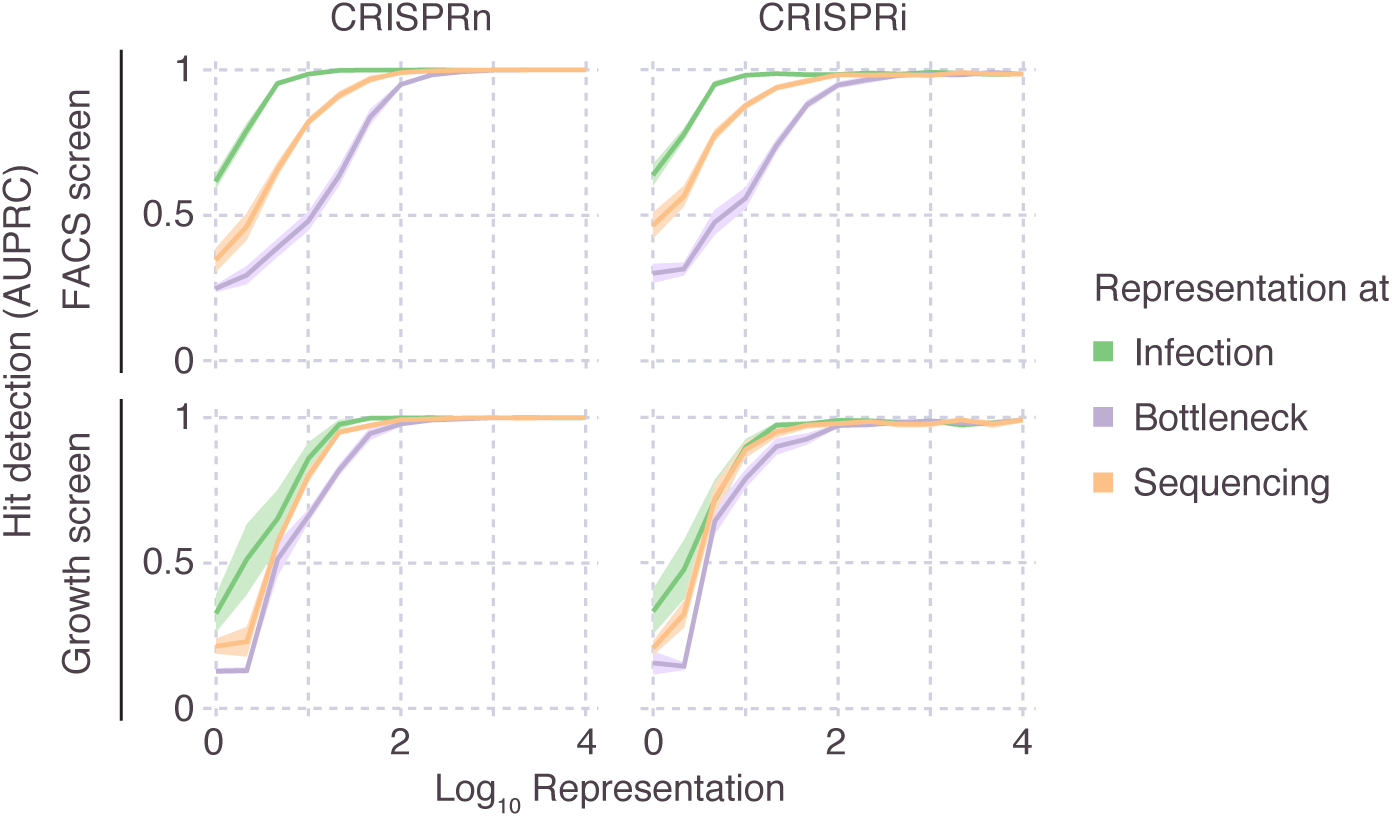
Importance of representation of library elements at different stages of the screen. CRISPulator simulations reveal the effect of library representation at different screen stages (Transfection, bottlenecks, sequencing) on hit detection. Simulations were run for FACS-based screens (top row) and growth-based screens (bottom row). Lines and light margins represent means and 95% confidence intervals, respectively, for 10 independent simulation runs.

For FACS screens using a given number of cells, an important decision is how extreme the cutoffs defining the “high-reporter” and “low-reporter” bins should be. CRISPulator simulation suggests that separating and comparing the cells with the top quartile and bottom quartile reporter activity results in the optimal detection of hit genes (**Fig. 9**). Closer inspection revealed that while both signal (sgRNA frequency differences between the two populations) and the noise (due to lower representation in the sorted population) decrease with larger bin sizes, the signal-to-noise ratio reaches a local maximum around 25% (**Fig. 10**), close to the bin size chosen fortuitously in published studies [8, 9].

**Figure 9.**
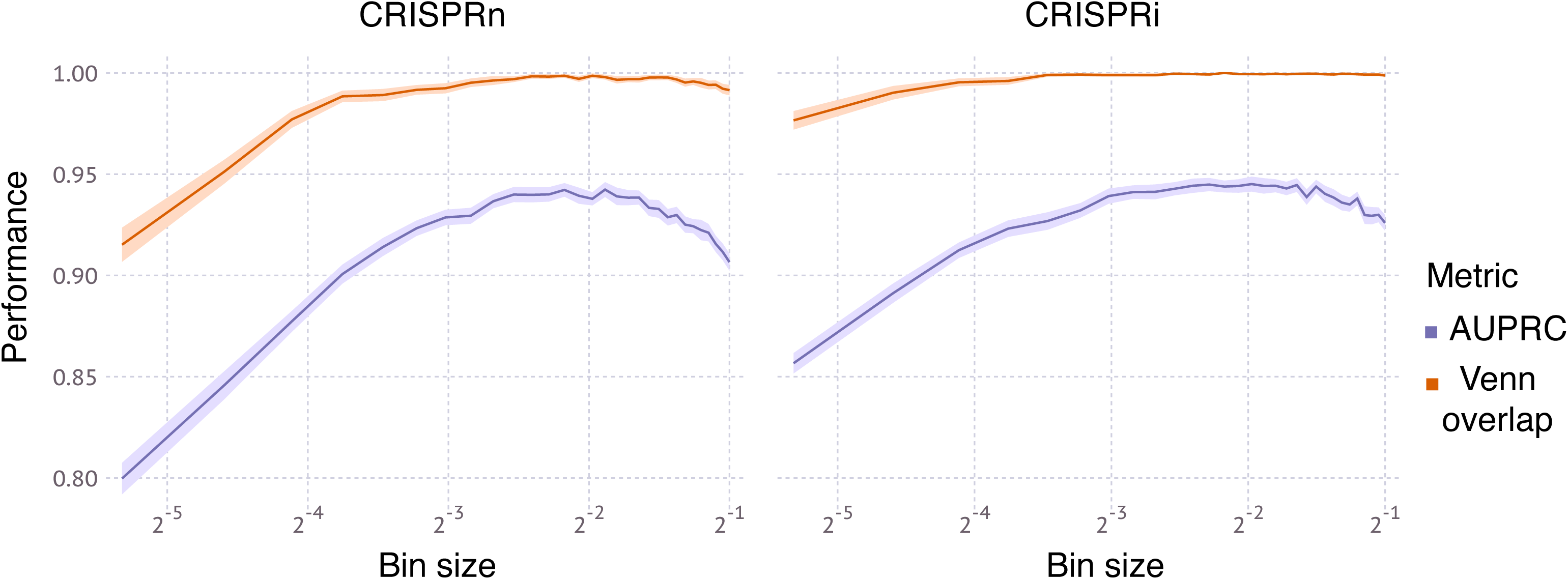
Effect of bin size on performance of FACS-based screens. Simulations were run for 100x representation at the transfection, bottleneck and sequencing stages. Lines and light margins represent means and 99% confidence intervals, respectively, for 100 independent simulation runs.

**Figure 10.**
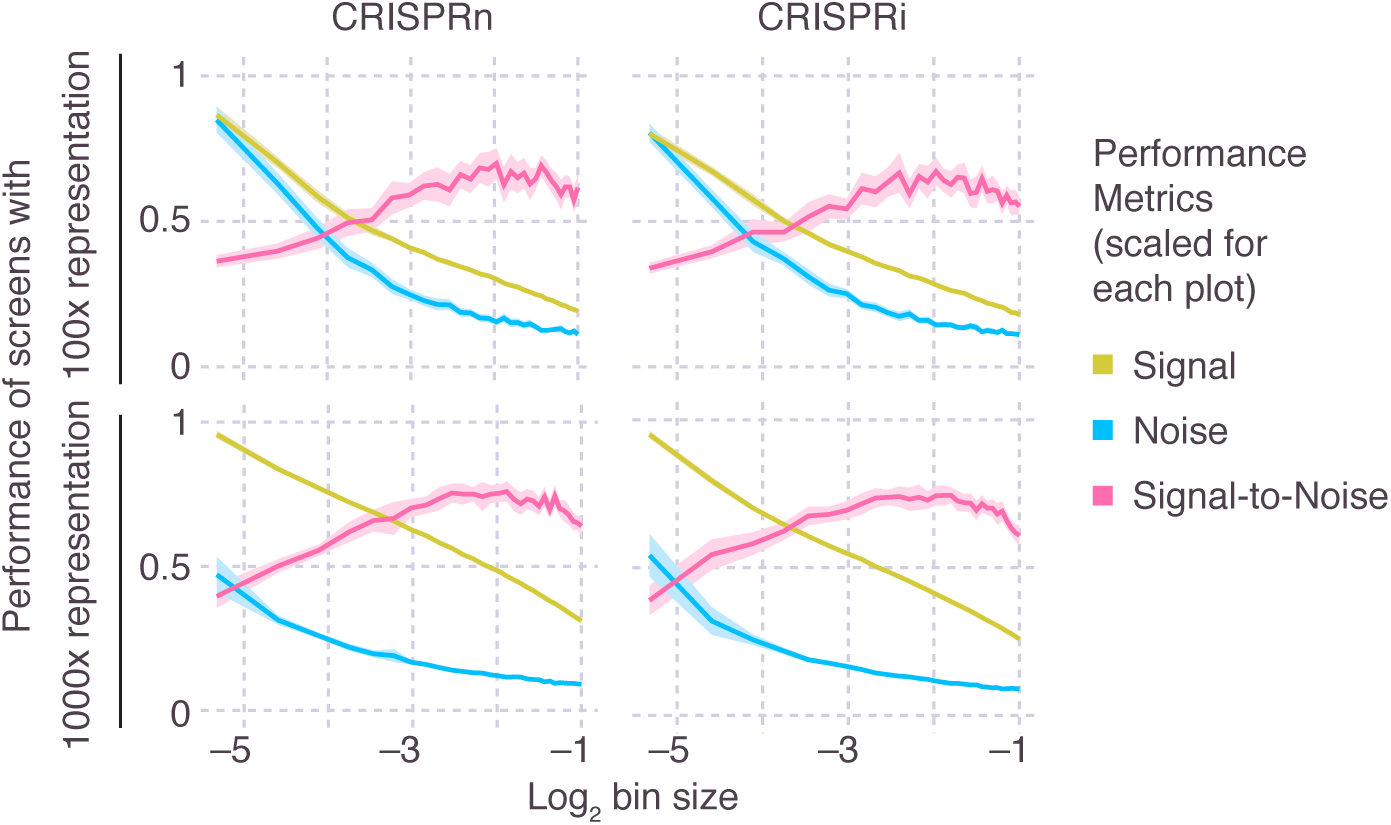
Effect of bin size on signal and noise of FACS-based screens. For FACS-based screens, the effect of the size of the sorted bins (see Fig. 1) on metrics for signal, noise, and signal-to-noise ratio (scaled within each plot) is shown. Metrics are defined in the Online Methods. Simulations were run for 10x representation (top row) or 100x representation (bottom row) at the transfection, bottleneck and sequencing stages. Lines and light margins represent means and 99% confidence intervals, respectively, for 25 independent simulation runs.

For growth-based screens, the duration of the screen influences the signal (by amplifying differences in frequency due to different growth phenotypes) but also the noise (by increasing the number of Poisson sampling bottlenecks generated by cell passaging or repeated applications of selective pressure). Interestingly, CRISPulator suggests that the effect of screen duration on optimal performance is different for genes with positive and negative phenotypes, and strongly depends on the presence of genes with positive phenotypes (**Fig. 11**). While genes with positive phenotypes (increased growth / survival) were detected more reliably after longer screens, genes with negative phenotypes (decreased growth / survival) were optimally detected in screens of intermediate duration, and their detection in longer screens rapidly declined if genes with stronger positive phenotypes were present in the simulated genome. While genes with positive phenotypes are rare in screens based on growth in standard conditions [5-7], selective pressures, such as growth in the presence of toxin, can reveal strong positive phenotypes for genes conferring resistance to the selective pressure [7]. The optimal screen length for growth-based screens was dictated by a local maximum of the signal-to-noise ratio, which itself depended on the representation: screens with lower representation were performing better at shorter duration (**Fig. 12**). Our results therefore predict that especially for growth-based screens using selective pressures, and screens implemented with low representation, short durations are preferable.

**Figure 11.**
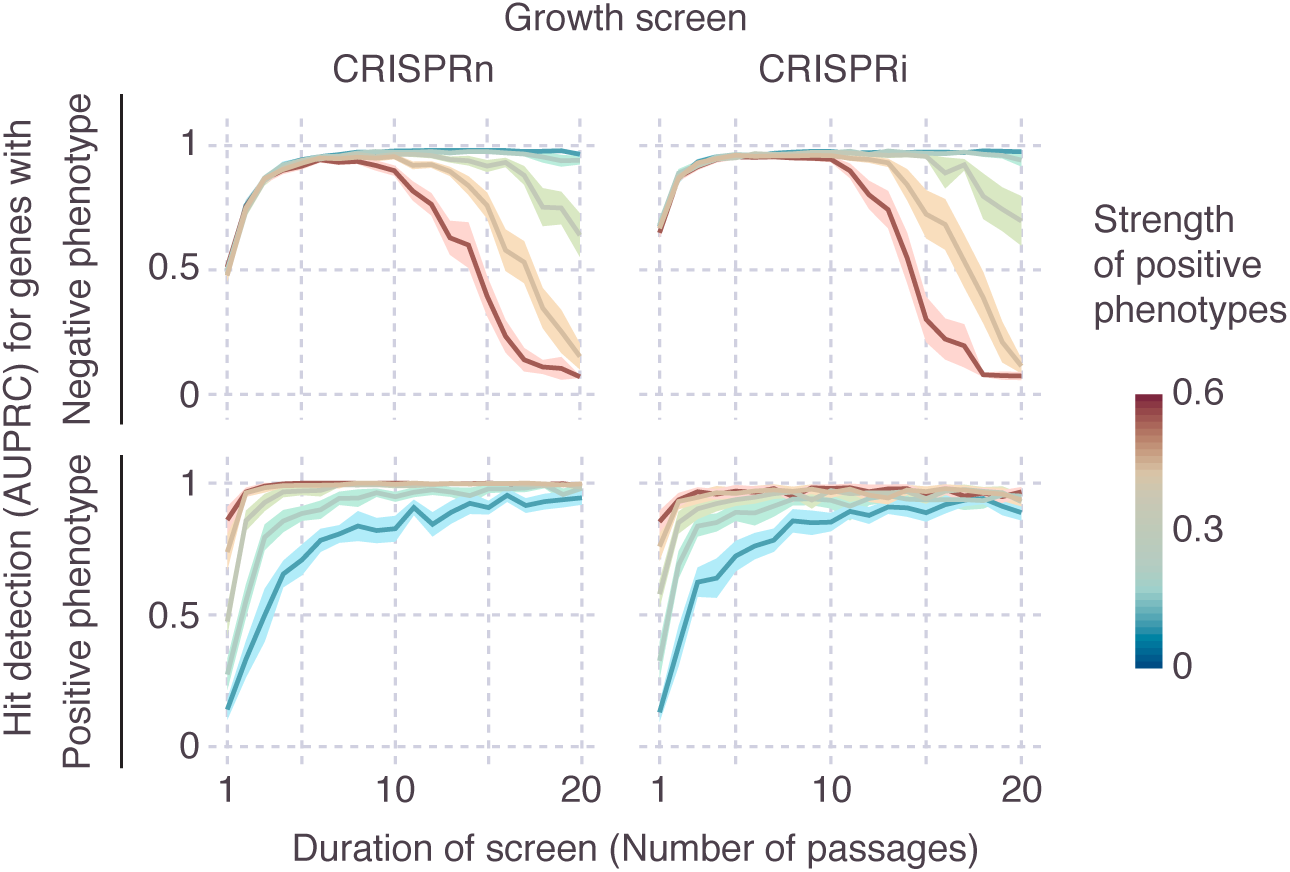
Effect of positive phenotypes on growth-based screens. For growth-based screens, the presence of genes with positive phenotypes (fitter than wild type) strongly influences hit detection as a function of screen duration. Screens were simulated for a set of genes in which 10% of all genes had negative phenotypes (less fit than wild type), and 2% of genes had positive phenotypes. The strength of positive phenotypes was varied, as encoded by the heat map. Hit detection was quantified separately for genes with negative phenotypes (top row) and genes with positive phenotypes (bottom row). Simulations were carried out for screens with different durations, as measured by the number of passages. Lines and light margins represent means and 95% confidence intervals, respectively, for 25 independent simulation runs. In **a** and **c**, hit detection is measured as Area under the Precision-Recall curve (AUPRC), as detailed in the Online Methods.

**Figure 12.**
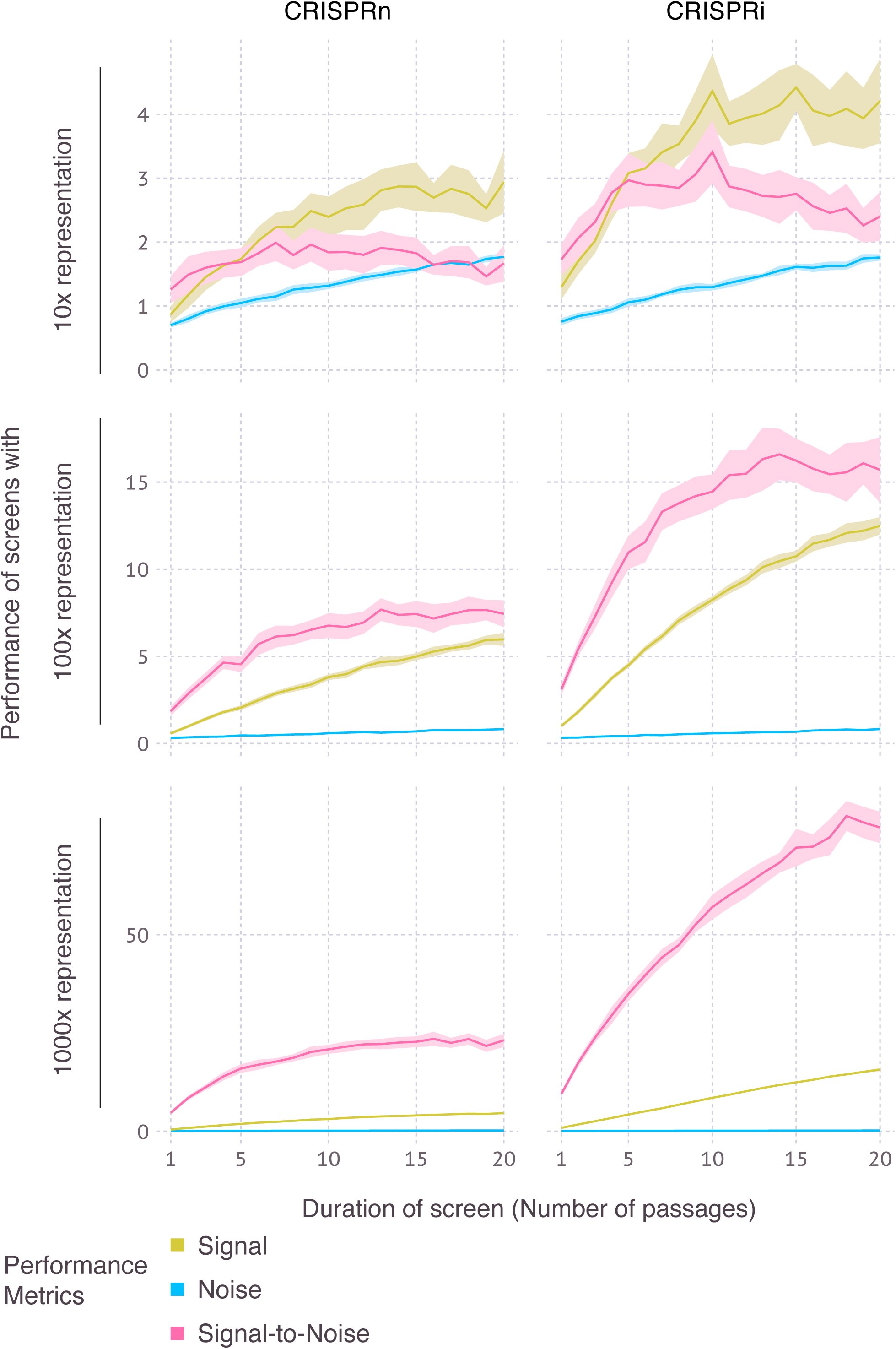
Effect of duration of growth-based screens on performance. Screens were simulated for a set of genes in which 10% of all genes had negative phenotypes (less fit than wild type). Simulations were carried out for screens with different durations, as measured by the number of passages, and for different representations at the transfection, bottleneck and sequencing stages. Metrics for signal, noise, and signal-to-noise ratio are defined in the Online Methods. Lines and light margins represent means and 95% confidence intervals, respectively, for 25 independent simulation runs.

While CRISPRn and CRISPRi screens performed similarly in the simulations described above (**Fig. 8-11**), separate evaluation of genes with linear versus sigmoidal phenotype-knockdown relationship revealed that CRISPRn outperforms CRISPRi for the detection of sigmoidal genes (which require very stringent knockdown to result in a phenotype), whereas CRISPRi performs relatively better for genes with a linear knockdown-phenotype relationship (**Fig. 13**).

**Figure 13.**
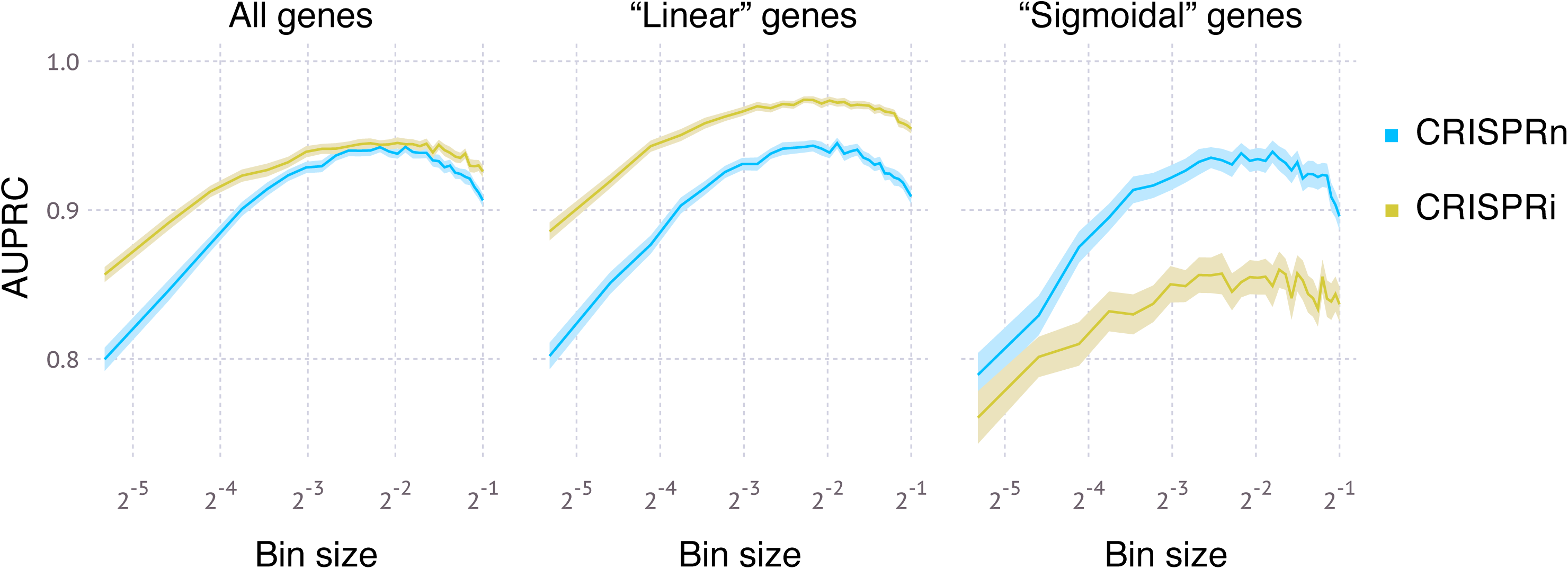
Comparison of CRISPRn and CRISPRi screen performance for genes with different knockdown-phenotype relationships. Simulations of FACS-based screens were run for 100x representation at the transfection, bottleneck and sequencing stages. The simulated genome contained 75% of genes with a linear knockdown-phenotype relationship and 25% of genes with a sigmoidal knockdown-phenotype relationship, as defined in the Online Methods. Performance in hit detection was quantified as AUPRC either for all genes, or only for linear or sigmoidal genes. Lines and light margins represent means and 99% confidence intervals, respectively, for 100 independent simulation runs.

## Discussion

CRISPulator revealed several non-obvious rules for the design of pooled genetic screens, illustrating its usefulness. Since certain parameters used by CRISPulator (such as the quality of sgRNA libraries or the signal-to-noise of FACS-based phenotypes) are estimates informed by published data, but not directly known, the predicted screen performance does not represent absolute performance metrics. Rather, the goal is to predict the relative performance of screens conducted with different experimental parameters to enable researchers to optimize those parameters. The simulated sequencing reads generated by CRISPulator (**Fig. 10**) recapitulate patterns observed in experimental data (**Fig. 14**), thereby facilitating the interpretation of suboptimal experimental data and providing a tool to predict which experimental parameters need to be changed to obtain data more suitable for robust hit detection

**Figure 14.**
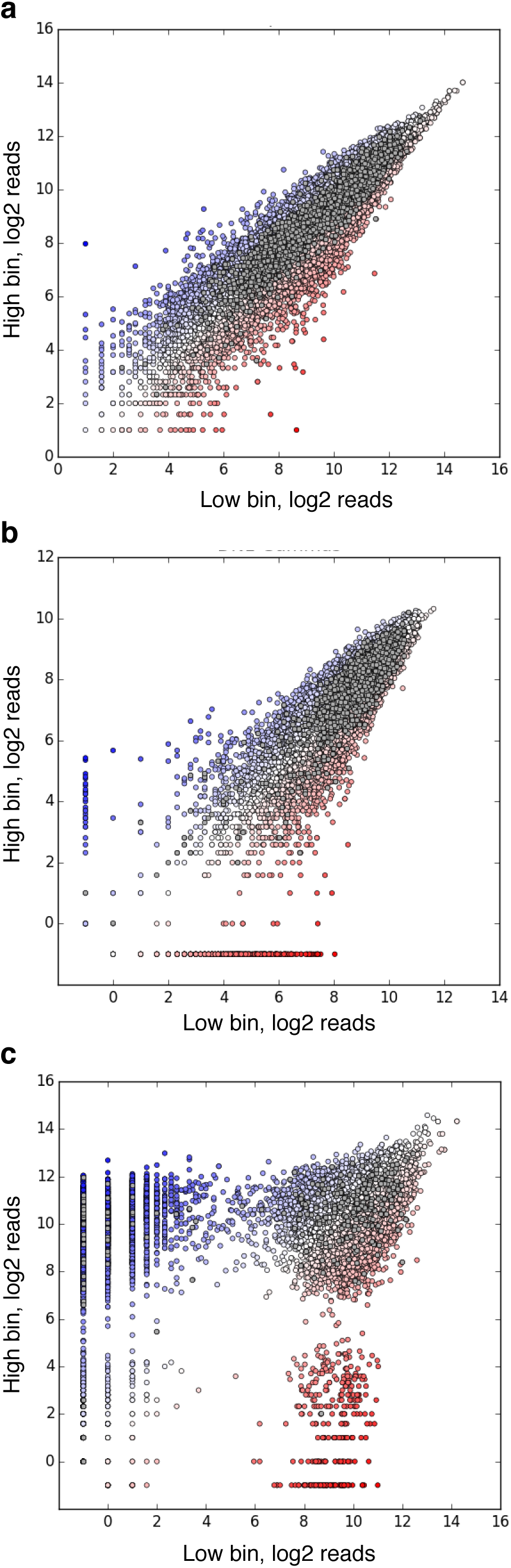
Experimental data from FACS-based screens resembles simulated data shown in Figure 6. Grey dots: non-targeting sgRNAs, dots on a red-white-blue color scale: targeting sgRNAs. Number of deep sequencing reads for each sgRNA in two populations separated based on a fluorescent reporter signal are shown. (**a**) Screen carried out with high representation at all stages. (**b**) Screen with low representation at the infection stage. (**c**) Screen with low representation at the selection stage.

## Conclusions

CRISPulator facilitates the design of pooled genetic screens by enabling the exploration of a large space of experimental parameters *in silico*, rather than through costly experimental trial and error. For pooled genetic screens in animal models, such as mice, choices of experimental parameters can also have ethical implications, namely the numbers of animals required to power the study. As larger numbers of pooled genetic screens are published, we will further refine the assumptions underlying the simulation using empirical data.

## Methods

### Code implementation and availability

CRISPulator was implemented in Julia (http://julialang.org), a high-level, high-performance language for technical computing. We have released the simulation code as a Julia package, Crispulator.jl. The software is platform-independent and is tested on Linux, OS X (macOS), and Windows. Installation details, documentation, source code, and examples are all publicly available at http://crispulator.ucsf.edu.

### Simulated genome

A genome is defined by assigning a numerical, “true” phenotype to a number of genes. All of our results featured here used 500 genes in each simulation. 75% of genes were assigned a phenotype of 0 (wild-type), and 5% of genes were modeled as negative control genes, also with a phenotype of 0. 10% of genes were assigned a positive phenotype randomly drawn (unless otherwise indicated) from a Gaussian distribution with μ=0.55 and σ=0.2 (clamped between [0.1, 1.0]), and 10% of genes were assigned a negative phenotype randomly drawn from an identical distribution except with μ=-0.55 and clamping [-1.0, -0.1] (**Fig. 2**). Next, each gene was randomly assigned a phenotype-knockdown function (**Fig. 3**) to simulate different responses of genes to varying levels of knockdown. 75% of genes were assigned a linear function that linearly interpolates between 0 and the “true” phenotype from above as a function of knockdown, the remaining 25% of genes were assigned a sigmoidal function with an inflection point, *p*, drawn from a distribution with a mean of 0.8 and standard deviation of 0.2; the width of the inflection region, *k*, (over which a phenotype increased from 0 to the “true” phenotype, *l*) was drawn from a normal distribution with a mean of 0.1 and a standard deviation of 0.05. The function *f* was defined as follows: 
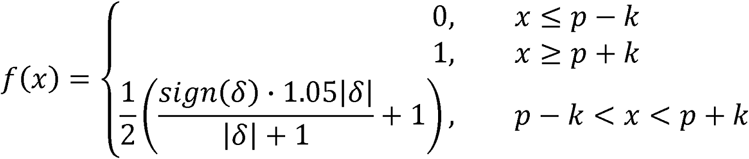
where 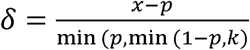

This specific sigmoidal function was chosen over the more standard special case of the logistic function or the Gompertz function because it is highly tunable and has a range between 0 and *1* on a domain of [0, 1].

### Simulated sgRNA libraries

CRISPRn and CRISPRi sgRNA libraries are generated to target the simulated genome. For the results featured here, each gene was targeted by 5 independent sgRNAs. For CRISPRi screens, each sgRNA was randomly assigned a knockdown efficiency from a bimodal distribution (**Fig. 4**): 10% of sgRNAs had low activity with a knockdown drawn from a Gaussian (μ=0.05, σ=0.07), 90% of guides had high activity drawn from a Gaussian (μ=0.90, σ=0.1). We assumed such a high rate of active sgRNAs based on our recently developed highly active CRISPRi sgRNA libraries [10]. For CRISPRn screens, high-quality guides all had a maximal knockdown efficiency of 1.0 and were 90% of the population (the 10% low-activity CRISPRn guides were drawn from the same Gaussian (μ=0.05, σ=0.07) as above). The initial frequency distribution of sgRNAs in the library was modeled as a log-normal distribution such that a guide in the 95^th^ percentile of frequencies is 10 times as frequent as one in the 5^th^ percentile (**Fig. 5**), which is typical of high-quality libraries in our hands [7].

### Simulated screens

All steps of the pooled screens are simulated discretely. Infections are modeled as a Poisson process with λ=M.O.I of the infection. The initial pool of cells is randomly infected by sgRNAs based on the frequency of each sgRNA in the library. A λ=0.25 is used unless otherwise noted, which is commonly used to approximate single-copy infection [11]. Only cells with a single sgRNA are then used in subsequent steps, which is P(x= 1; Poisson(λ= 0.25)) ≈ 19.5% of the initial pool.

For CRISPRi screens, phenotypes for each cell were determined based on the sgRNA knockdown efficiency (from above) and based on both the phenotype and the knockdown-phenotype relationship of the targeted gene. For CRISPRn screens, phenotypes for each cell were set using using sgRNA knockdown efficiency (specific for CRISPRn screens, see previous paragraph) and the gene phenotype. If a cell was infected with a low-quality CRISPRn guide, it behaved similarly to a low-quality CRISPRi guide, i.e. very close to no activity. All cells with high-quality guides CRISPRn guides had a 1/9, 4/9, or 4/9 chance of having 0%, 50%, or 100% knockdown efficiency, respectively. This knockdown efficiency was then used with the knockdown-phenotype relationship and true phenotype of the gene to calculate the observed phenotype. The assumption that only bi-allelic frame-shift mutations lead to a phenotype in CRISPRn screens for most sgRNAs is supported by the empirical finding that in-frame deletions mostly do not show strong phenotypes, unless they occur in regions encoding conserved residues or domains [10]. To mitigate this issue, some CRISPRn screens have been conducted in quasi-haploid cell lines [6].

FACS sorting was simulated by convoluting the theoretical phenotypes of each cell independently with a Gaussian (μ= 0, σ) where σ is a tunable “noise” parameter, reflecting biological variance in fluorescence intensity of isogenic cells. The number of cells prior to this step is termed the bottleneck representation and is tunable. Post-convolution, cells were sorted according to their new, “observed” phenotype and then the bottom X percentile and top X percentile (X was real value between 0 and 50) were taken as the two comparison bins.

Growth experiments were simulated as follows: (1) in the time frame that WT cells (true phenotype=0) divide once, cells with the maximal negative phenotype, -1, do not divide, and cells with maximal positive phenotype divide twice. For cells with phenotypes in between 0 and ±1, cells randomly pick whether they behave like WT cells or maximal phenotype cells weighted by their phenotype (i.e. cells with phenotypes close to 0 behave mostly like WT cells). (2) After one timestep where WT cells double once, a random subsample of the cells is taken. The size of the bottleneck is tunable. (3) This is repeated *n* number of times. Finally, the samples of cells at t=0 and t=n are taken as the two populations for comparison.

Sample preparation was simulated by taking the frequencies of each guide in the cells after selection and constructing a categorical distribution with the frequencies as the weights. Next-generation sequencing was then simulated by sampling from this categorical distribution up to the number of total reads.

### Evaluation of screen performance

Based on the simulated sequencing read counts, P values and gene-level phenotypes were calculated for each gene essentially as previously described [3, 7]. Briefly, observed sgRNA phenotypes were calculated as log_2_ ratios of sgRNA frequencies in two cell populations. Gene-level phenotypes were calculated by averaging the sgRNA phenotypes. P values were calculated based on the Mann-Whitney rank-sum test by comparing the phenotypes of sgRNAs targeting a given gene with the phenotypes of negative control sgRNAs. Genes were ranked by the product of the absolute gene-level phenotype and their – log_10_ P value to call hit genes. Screen performance was quantified in two ways: As the overlap of the top 50 called hit genes with the top 50 actual hit genes (based on true phenotype), or as the area under the precision-recall curve (AUPRC). AUPRC was chosen over the more common area under the receiver operator characteristic (AUROC) due to the highly skewed nature of the generated dataset (<20% of dataset is made up of true hits). The AUPRC was calculated using a lower trapezoidal estimator, which had been previously shown to be a robust estimator of the metric [12]. For Figures 10 and 12, the “signal” of an experiment was defined as the median signal for true hit genes (ones initially labeled as having a positive or negative phenotype). The true hit gene signal was calculated as the average ratio of the log_2_ fold change over the theoretical phenotype of all guides targeting that gene. Guides that dropped out of the analysis were excluded from the signal calculation. “Noise” was quantified as the standard deviation of negative-control sgRNA phenotypes, and the “signal-to-noise” ratio was the ratio of these two metrics. For display purposes, all are normalized in each graph.

## List of abbreviations

AUPRC: area under the precision-recall curve
CRISPRi: CRISPR interference
CRISPRn: CRISPR nuclease
FACS: fluorescence-activated cell sorting
sgRNA: single guide RNA

## Declarations

### Ethics approval and consent to participate

Not applicable.

### Consent for publication

Not applicable.

### Availability of data and materials

The software, CRISPulator, described in this study, which was also used to generate the data, is publicly available at http://crispulator.ucsf.edu (an archived version is available at https://doi.org/10.5281/zenodo.345188). It is released under Apache License 2.0 and there are no additional restrictions for commercial use.

### Authors’ contributions

TN and MK designed and analyzed the research and wrote the manuscript. TN developed the CRISPulator software.

## Acknowledgments

We thank Ruilin Tian and Daniel Asarnow for feedback on this manuscript, and Diane Nathaniel, John Chen, Kathleen Keough and Xiaoyan Guo for sharing unpublished experimental data.

## References

1. Kaelin WG, Jr.: Molecular biology. Use and abuse of RNAi to study mammalian gene function. Science 2012, 337: 421–422.

2. Kampmann M, Horlbeck MA, Chen Y, Tsai JC, Bassik MC, Gilbert LA, Villalta JE, Kwon SC, Chang H, Kim VN, Weissman JS: Next-generation libraries for robust RNA interference-based genome-wide screens. Proc Natl Acad Sci U S A 2015, 112:E3384–E3391.

3. Kampmann M, Bassik MC, Weissman JS: Integrated platform for genome-wide screening and construction of high-density genetic interaction maps in mammalian cells. Proc Natl Acad Sci US A 2013, 110:E2317–2326.

4. Shalem O, Sanjana NE, Zhang F: High-throughput functional genomics using CRISPR-Cas9. Nat Rev Genet 2015, 16: 299–311.

5. Shalem O, Sanjana NE, Hartenian E, Shi X, Scott DA, Mikkelsen TS, Heckl D, Ebert BL, Root DE, Doench JG, Zhang F: Genome-scale CRISPR-Cas9 knockout screening in human cells. Science 2014, 343: 84–87.

6. Wang T, Wei JJ, Sabatini DM, Lander ES: Genetic screens in human cells using the CRISPR-Cas9 system. Science 2014, 343: 80–84.

7. Gilbert LA, Horlbeck MA, Adamson B, Villalta JE, Chen Y, Whitehead EH, Guimaraes C, Panning B, Ploegh HL, Bassik MC, et al: Genome-Scale CRISPR-Mediated Control of Gene Repression and Activation. Cell 2014, 159: 647–661.

8. Sidrauski C, Tsai JC, Kampmann M, Hearn BR, Vedantham P, Jaishankar P, Sokabe M, Mendez AS, Newton BW, Tang EL, et al: Pharmacological dimerization and activation of the exchange factor eIF2B antagonizes the integrated stress response. Elife 2015, 4: e07314.

9. DeJesus R, Moretti F, McAllister G, Wang Z, Bergman P, Liu S, Frias E, Alford J, Reece-Hoyes JS, Lindeman A, et al: Functional CRISPR screening identifies the ufmylation pathway as a regulator of SQSTM1/p62. Elife 2016, 5.

10. Horlbeck MA, Gilbert LA, Villalta JE, Adamson B, Pak RA, Chen Y, Fields AP, Park CY, Corn JE, Kampmann M, Weissman JS: Compact and highly active next-generation libraries for CRISPR-mediated gene repression and activation. Elife 2016, 5.

11. Fellmann C, Zuber J, McJunkin K, Chang K, Malone CD, Dickins RA, Xu Q, Hengartner MO, Elledge SJ, Hannon GJ, Lowe SW: Functional identification of optimized RNAi triggers using a massively parallel sensor assay. Mol Cell 2011, 41: 733–746.

12. Boyd K, Eng KH, Page CD: In Machine Learning and Knowledge Discovery in Databases. Edited by Blockeel H, Kersting K, Nijssen S: Springer; 2013: 451–466

